# Visualizing alpha-synuclein and iron deposition in M83 mouse model of Parkinson’s disease *in vivo*

**DOI:** 10.1101/2023.06.28.546962

**Authors:** Nadja Straumann, Benjamin F. Combes, Xose Luis Dean Ben, Rebecca Sternke-Hoffmann, Juan A. Gerez, Ines Dias, Zhenyue Chen, Benjamin Watts, Iman Rostami, Kuangyu Shi, Axel Rominger, Christian R Baumann, Jinghui Luo, Daniela Noain, Roger M. Nitsch, Nobuyuki Okamura, Daniel Razansky, Ruiqing Ni

**Author notes:** Corresponding author: Ruiqing Ni, Address: Wagistrasse 12, 8952 Zurich, Switzerland.

## Abstract

**Background:** Abnormal alpha-synuclein and iron accumulation in the brain play an important role in Parkinson’s disease (PD). Herein, we aim at visualizing alpha-synuclein inclusions and iron deposition in the brains of M83 (A53T) mouse models of PD *in vivo*.

**Methods:** Fluorescently labelled pyrimidoindole-derivative THK-565 was characterized by using recombinant fibrils and brains from 10-11 months old M83 mice, which subsequently underwent *in vivo* concurrent wide-field fluorescence and volumetric multispectral optoacoustic tomography (vMSOT) imaging. The *in vivo* results were verified against structural and susceptibility weighted imaging (SWI) magnetic resonance imaging (MRI) at 9.4 Tesla and scanning transmission X-ray microscopy (STXM) of perfused brains. Brain slice immunofluorescence and Prussian blue staining were further performed to validate the detection of alpha-synuclein inclusions and iron deposition in the brain, respectively.

**Results:** THK-565 showed increased fluorescence upon binding to recombinant alpha-synuclein fibrils and alpha-synuclein inclusions in post-mortem brain slices from patients with Parkinson’s disease and M83 mice. *i.v.* administration of THK-565 in M83 mice showed higher cerebral retention at 20 and 40 minutes post-injection by wide-field fluorescence compared to non-transgenic littermate mice, in congruence with the vMSOT findings. SWI/phase images and Prussian blue indicated the accumulation of iron deposits in the brains of M83 mice, presumably in the Fe^3+^ form, as evinced by the STXM results.

**Conclusion:** We demonstrated *in vivo* mapping of alpha-synuclein by means of non-invasive epifluorescence and vMSOT imaging assisted with a targeted THK-565 label and SWI/STXM identification of iron deposits in M83 mouse brains *ex vivo*.

## 1. Introduction

Alpha-synucleinopathies such as Parkinson’s disease (PD), dementia with Lewy bodies (DLB), and multiple system atrophy (MSA) are some of the most common neurodegenerative diseases [1]. Neuronal inclusions formed by β-sheet rich fibrillar alpha-synuclein (αSyn) termed Lewy bodies, pale bodies, and Lewy neurites are pathological hallmarks of PD. The accumulation of misfolded αSyn inclusions precedes the loss of dopaminergic neurons in the substantia nigra and is an important target for early diagnosis [1]. Iron is linked to important biological processes in the brain, such as oxygen transportation and neurotransmitter synthesis [2]. For iron deposit detection, *in vivo* magnetic resonance imaging (MRI), such as using T_2_* [3, 4], susceptibility weighted imaging (SWI) and quantitative susceptibility mapping (QSM) [5, 6], has demonstrated brain iron accumulation in patients with atypical PD syndromes [7] and PD [8], as well as in animal models of PD [3]. Abnormal accumulation of iron has been shown to confer neurotoxicity to nigral neurons [9], induce dopaminergic damage [10], and promote αSyn aggregation. In turn, αSyn aggregation disrupts iron metabolism, leading to elevated iron accumulation and redistribution within neurons and promoting ferroptosis in animal models [11, 12].

A few chemical structures and PET imaging ligands targeting αSyn have been identified and evaluated *in vitro* or in animal models (rodents, minipigs, and non-human primates)[13], such as [^18^F]C05-05, [^11^C]MODAG-001, [^11^C]MK-7337, [^18^F]4FBox, [^18^F]AS69 affibody and antibody-based [^124^I]RmAbSynO2-scFv8D3 [14–19]. In addition, fluorescence-emitting imaging probes have been developed and evaluated in post-mortem human brain tissue with αSyn inclusions [20, 21]. *In vivo* fluorescence imaging of αSyn inclusions in animal models has been reported only using rats injected with labelled fluorescence atto-647 αSyn fibrils [22] or in the retinas of mice overexpressing αSyn fused to green fluorescent protein [23]. On the other hand, fluorescence imaging approaches such as 2-photon microscopy and diffuse optical imaging have a limited penetration depth. MRI using ScFv-conjugated superparamagnetic iron oxide nanoparticles W20-SPIONs in A53T mice [24] provided whole brain coverage despite relatively low sensitivity. Volumetric multispectral optoacoustic tomography (vMSOT) has the unique features of high sensitivity of optical contrast and a high spatial resolution of ∼110 µm by ultrasound [25, 26] and can cover a penetration depth of several cm sufficient for mouse brain imaging. This approach provides detection of cerebral hemodynamics based on deoxy-/oxyhemoglobin (Hb/HbO_2_) contrasts, as well extrinsic contrast agents after unmixing, such as neuroinflammation, Aβ and tau deposits in the brain of animal models of diseases [27, 28].

In this study, we aimed to visualize αSyn and iron deposits in the brains of a transgenic mouse model of PD, the M83 (A53T) line, using concurrent epifluorescence-vMSOT, high-field MRI, and scanning transmission X-ray microscopy. We utilized the novel pyrimidoindole derivative THK-565 for *in vivo* imaging of αSyn inclusions.

## 2. Methods

### 2.1 Post-mortem human brain tissue

One PD case (male, 56 years old) with a clinical diagnosis confirmed by pathological examination of Lewy bodies (Braak LB 5, without tau and Aβ) was included in this study. Paraffin-embedded autopsy brain tissues from the medulla oblongata with high αSyn inclusion accumulation were obtained from the Netherlands Brain Bank (NBB), Netherlands. All materials had been collected from donors or from whom written informed consent for a brain autopsy and the use of the materials and clinical information for research purposes had been obtained by the NBB. The study was conducted according to the principles of the Declaration of Helsinki and subsequent revisions. All experiments on autopsied human brain tissue were carried out in accordance with ethical permission obtained from the regional human ethics committee in Canton Zurich and the medical ethics committee of the VU Medical Center for the NBB tissue.

### 2.2 Animal models

Transgenic mice carrying the A53T-mutated human αSyn gene [29] (M83, both sexes, n = 15) and non-transgenic littermates (NTLs, both sexes, n = 9) were used. Animals were housed in individually ventilated cages inside a temperature-controlled room under a non-inverted 12-hour dark/light cycle. Two arcAβ transgenic mice [28, 30] overexpressing the human APP695 transgene containing the Swedish (*K670N/M671L*) and Arctic (*E693G*) mutations under the control of the prion protein promoter and two age-matched NTL of both sexes (18 months of age) were used. Two *MAPT* P301L transgenic mice overexpressing human 2N/4R tau under the neuron-specific Thy1.2 promoter (pR5 line, C57B6. Dg background) [31–33], and two respective NTL mice were used. Pelleted food (3437PXL15, CARGILL) and water were provided ad libitum. All experiments were performed in accordance with the Swiss Federal Act on Animal Protection and were approved by the Cantonal Veterinary Office Zurich (permit numbers: ZH024/21, ZH162/20).

### 2.3 Chemicals

8-Methyl-10,10-dimethzl-10a-[4-[4-(dimethzlamino)phenyl-1,3,butadienyl]-3,4,10,10 atetrahyro-pyrimido[1,2,a] indol-2(1H)-one (THK-565) was synthesized and kindly provided by Prof. Nobuyuki Okamura (Tohoku University, Japan) (MW 401.56, chemical structure shown in **Fig. 1a**). Other chemicals and reagents were commercially purchased (details in **Suppl. Table 1**).

**Fig 1.**
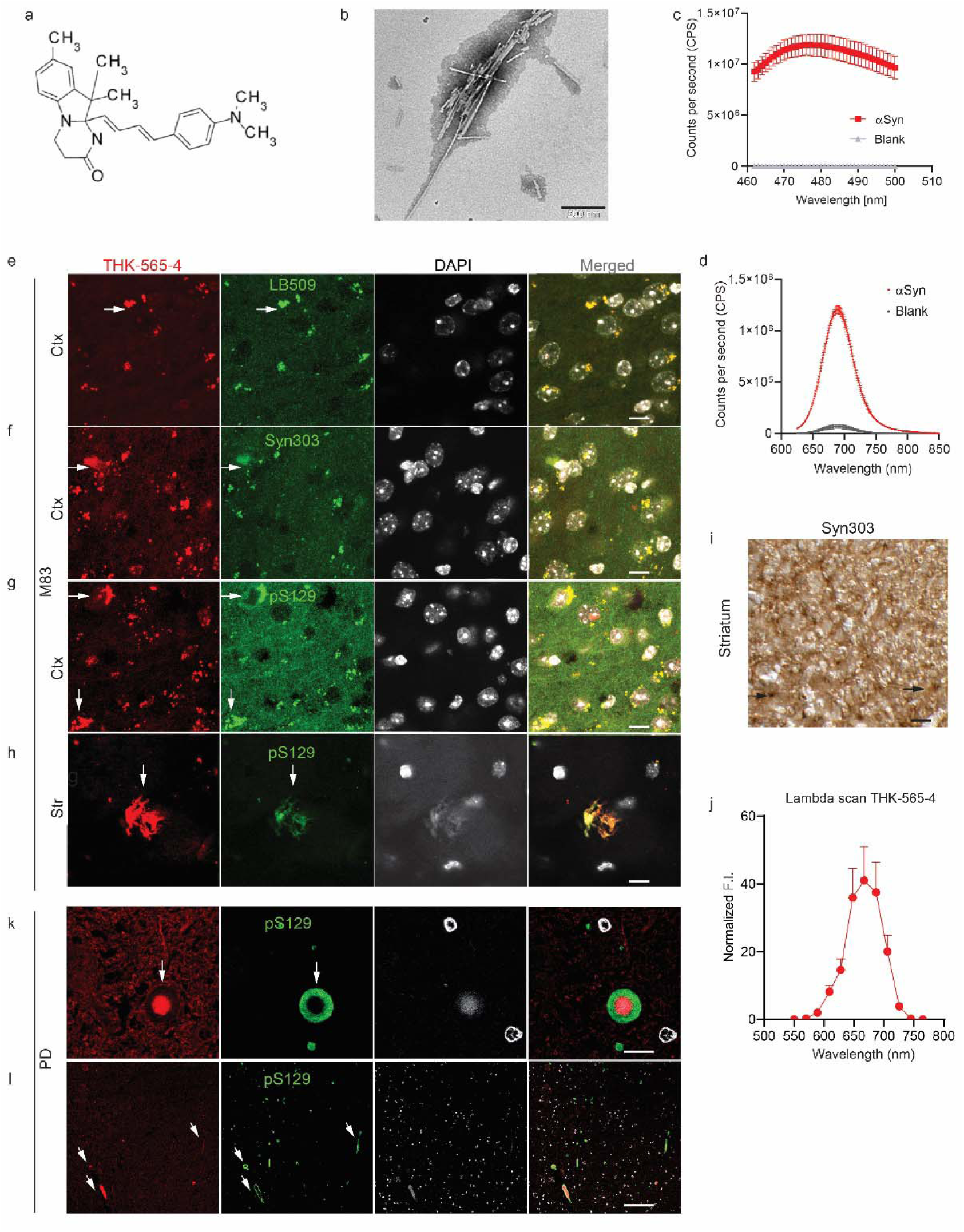
Characterization of THK-565 in recombinant αSyn fibrils and M83 mouse and post-mortem PD brain. (**a**) Chemical structure of THK-565; (**b**) Transmission electron microscopy characterization of recombinant αSyn fibril; Scale bar = 200 nm; (**c**) Thioflavin T assay of recombinant αSyn (red) fibril and blank (gray); (**d**) Spectrofluorometric measurements of the binding of THK-565 to recombinant αSyn (red) fibril and blank (gray, dd. water); (**e-h**) Immunofluorescence staining using THK-565 (red), with anti-αSyn antibodies LB509, Syn303, anti-p-αSyn antibody pS129 (green) on cortex (Ctx) and striatum (Str) of M83 mouse brain; (**i**) Immunocytochemistry using Syn303 antibodies on M83 mouse Str; arrow indicates αSyn inclusions. (**j**) Lambda scan of THK-565-stained αSyn inclusions in the M83 mouse brain. (**k, l**) Immunofluorescence staining using THK-565 (red), with pS129 (green) on medulla oblongata of postmortem tissue from patient with PD; nuclei were counterstained using DAPI (gray). Scale bar = 10 μm (e-h, j, k) and 50 μm (i, l);

### 2.4 *In vitro* fluorescence assay for the binding of ligands to recombinant Aβ_42_, K18 tau and αSyn fibrils

Recombinant Aβ_42_, K18 tau, and αSyn were expressed and produced by *E. coli* as described previously [34–36]. The fluorescent dyes were dissolved in Milli-Q H_2_O or dimethyl sulfoxide (DMSO) and further diluted in 1×PBS pH 7.4. The absorbance of the compounds was measured. Thioflavin T assays against Aβ_42_ and K18 tau and αSyn fibrils were performed as described previously [34, 36], with two independent experiments and three technical replicates (Fluoromax 4, Horiba Scientific, Japan). The dyes were then mixed with either 2 μL of αSyn, 5 μL of K18 tau (380 μg/ml) or 5 μL of Aβ_42_ fibril (80 μg/ml) solution in a 45 µL quartz cuvette (quartz SUPRASIL Ultra Micro Cell, Hellma). The solutions were incubated for 1 min at room temperature and resuspended, and fluorescence was measured with a spectrofluorometer (FluoroMax-4, Horiba Jobin Yvon, Japan) using a known excitation wavelength for the ligands.

### 2.5 *In vivo* and *ex vivo* imaging with the hybrid epifluorescence and vMSOT system

Concurrent epifluorescence imaging and vMSOT at pre, during, and post *i.v.* bolus injection of THK-565 was performed using a previously established hybrid system consisting of an epifluorescence fiberscope and a vMSOT imager [37, 38]. The field-of-view (FOV) has a diameter of 12 mm for epifluorescence imaging and 15×15×15 mm^3^ for vMSOT, hence covering the entire mouse brain. The spatial resolution is approximately 40 μm and 120 μm for epifluorescence and vMSOT, respectively [39, 40]. Mice were first anaesthetized with an initial dose of 4% isoflurane (Abbott, Cham, Switzerland) in an oxygen/air mixture (200/800 mL/min) and subsequently maintained at 1.5% isoflurane in oxygen/air (100/400 mL/min) throughout the measurement. The fur over the head of the mice was removed before they were placed in the prone position on a heating pad with feedback control to maintain a constant body temperature (PhysioSuit, Kent Scientific, USA). The mice were subsequently injected with a 10 μL bolus containing THK-565 (**Fig. 1a**, dissolved in dimethyl sulfoxide (DMSO), 0.1 M PBS pH 7.4 followed by 90 μL saline through the tail vein. A dose of 20 mg/kg body weight was chosen and used in the following experiment. For vMSOT, the pulse repetition frequency of the laser was set to 25 Hz, and the laser wavelength was tuned between 550 and 660 nm (5 nm step) on a per pulse basis. Epifluorescence imaging was performed by coupling the same beam from the pulsed optical parametric oscillator (OPO) laser into the excitation fibre bundle. The excited fluorescence signal was collected by an imaging fibre bundle comprised of 100,000 fibres and then projected onto an electron multiplying charge-coupled device (EMCCD) camera (Andor iXon Life 888, Oxford Instruments, UK). Dynamic signals from epifluorescence and vMSOT were recorded simultaneously and synchronized with an external device (Pulse Pal V2, Sanworks, USA). To determine the optimal imaging time window, one mouse was scanned every 30 min starting at 60 min post-injection of THK-565 until 320 min after the injection of THK-565. For the remaining mice, the scan time points were before injection (108 s duration), during injection (432 s duration with *i.v.* injection starting at 30 s after the beginning of acquisition) and 20, 40, 60 and 90 min post-injection (108 s duration each).

The reconstructed vMSOT images were spectrally processed to unmix the biodistribution of THK-565. For this, per-voxel least square fitting of the spectral signal profiles to a linear combination of the absorption spectra of HbO_2_ and THK-565 was performed [27, 37, 41]. Given that there is no brain atrophy in the mice (based on *ex vivo* MRI data described below), the resulting images were coregistered with a structural MRI atlas (Ma-Benveniste-Mirrione-T_2_) in PMOD 4.2 (Bruker, Germany) for volume-of-interest (VOI)-based data analysis (Bruker, Germany). The fluorescence and vMSOT signal intensity were adjusted by dose/weight and normalized to 0-1 scale.

### 2.6 *Ex vivo* MRI of the M83 mouse brain

M83 and NTLs were intracardially perfused under deep anaesthesia (ketamine/xylazine/acepromazine maleate (75/10/2 mg/kg body weight, i.p. bolus injection) with 0.1 M phosphate buffered saline (PBS, pH 7.4), followed by 4% paraformaldehyde in 0.1 M PBS (pH 7.4). The heads were post-fixed in 4% paraformaldehyde in 0.1 M PBS (pH 7.4) for 6 days and stored in 0.1 M PBS (pH 7.4) at 4 °C. Brains were not removed from the skull, which has been shown previously to preserve cortical and central brain structure. The heads were placed in a 15 ml centrifuge tube filled with perfluoropolyether (Fomblin Y, LVAC 16/6, average molecular weight 2700, SigmaLJAldrich, USA). MRI data were acquired on a BioSpec 94/30 with a cryogenic 2×2 radio frequency phased-array surface coil (overall coil size 20×27 mm^2^, Bruker BioSpin AG, Fällanden, Switzerland) with a coil system operating at 30LJK (Bruker BioSpin AG, Fällanden, Switzerland) for reception used in combination with a circularly polarized 86 mm volume resonator for transmission. For SWI, a global and MAPSHIM protocol with a field map (default settings) was used for shimming [42]. A 3D gradient-recalled echo SWI sequence was recorded with the following parameters: FOVLJ=LJ15×12×15 mm; image sizeLJ=LJ248×199×36 μm, resulting in a spatial resolutionLJ= 60×60LJ×438 µm. One echo with a TE =LJ12 ms; TR =LJ250 ms; flip angleLJ=LJ15°; number of averagesLJ=LJ4, acquisition scan timeLJ= 1 h 59 min 24 s. SW and phase images were computed using the SWI processing module in ParaVision 6.0.1 (Bruker, Ettlingen, Germany) with Gauss broadeningLJ=LJ1 mm and mask weightingLJ=LJ4. All SW images were compared with their phase image counterparts. MRI images were analysed in ITK SNAP [43] and evaluated by two people blinded to the genotype of the mice.

### 2.7 Histology, immunofluorescence and confocal imaging

We assessed the binding of THK-565 to αSyn inclusions in M83 mouse brains and post-mortem brain tissue from patients with PD, as well as to Aβ plaques and tau inclusions in arcAβ and P301L mouse brains, by using immunofluorescence staining. For mouse brain samples, the brain was cut into 40 μm-thick coronal sections using a vibratome (Leica VT1000S, Germany) for free-floating immunofluorescence immunohistochemistry or embedded in paraffin following routine procedures and cut into 5 μm-thick sections for histology. For paraffin-embedded post-mortem human brain tissue, 5 μm-thick sections were cut for histopathology. Costaining was performed using ligands and anti-phospho-tau antibody AT-8, anti-β-amyloid1-16 antibody 6E10, anti-αSyn antibodies LB509 (total αSyn), and Syn303 (oxidized αSyn), and pS129 (phosphorylated αSyn), as described earlier [44] (details in **Suppl. Table 1**). Counterstaining was performed using 4’,6-diamidino-2-phenylindole (DAPI).

Hematoxylin & eosin (HE) staining and Prussian blue (for iron deposit detection) staining were performed in M83 mouse brains. Brain sections were imaged at 20× magnification using an Axio Oberver Z1 and at 63× magnification using a Leica SP8 confocal microscope (Leica, Germany) for colocalization evaluations. A lambda scan using a Leica SP8 was performed on stained brain slices to further determine the fluorescent properties of THK-565. The images were analysed using Qupath and ImageJ (NIH, U.S.A).

### 2.8 STXM

X-ray spectromicroscopy (scanning transmission X-ray microscopy, STXM) was performed at the Swiss Light Source (SLS, Switzerland) at the PolLux beamline (X07DA). Samples were prepared by cutting adjacent brain sections of which Prussian blue staining was performed into 60 nm thin sections and placing on a copper EM grid. The samples were raster scanned with a focus beam using a Fresnel zone plate with a width of 40 nm, setting the spatial resolution limit of the measurement. The transmitted X-ray beam was detected by a scintillator and a photomultiplier tube. To map the distributions of iron species, paired images were taken at 705 to 730 eV.

### 2.9 Statistics

Group comparison of THK-565 absorbance in multiple brain regions at different time points was performed by using two-way ANOVA with Bonferroni *post hoc* analysis using GraphPad Prism 9 (GraphPad, USA). The difference in the fluorescence intensity acquired by epifluorescence imaging at different time points was compared using one-way ANOVA. All data are presented as the mean ± standard deviation. Pearson’s rank correlation analysis was used to compare vMSOT and epifluorescence imaging data and reliability analysis. Significance was set at ^∗^*p* < 0.05.

## 3 Results

### 3.1 *In vitro* fluorescence binding assays with recombinant fibrils and immunofluorescence in brain slices

First, we produced Aβ_42_, K18 tau, and αSyn fibrils using bacterially produced recombinant monomers and validated them using the ThT assay and TEM (**Figs. 1b, c; SFigs. 1a, b, d, e)**. THK-565 showed increased fluorescence emission in the presence of Aβ_42_, K18 tau and αSyn fibrils (**Fig. 1d, SFig. 1c, f**). The emission spectrum of THK-565 did not differentiate among the Aβ_42_, K18 tau and αSyn fibrils tested. THK-565 showed suitable absorbance and emission spectra for *in vivo* vMSOT.

Immunofluorescence staining of M83 mouse brain slices showed colocalization of the THK-565 signal with LB509-positive, Syn303-positive and pS129-positive αSyn inclusions (**Figs. 1e-h**). Immunocytochemistry using Syn303 antibodies further validated the presence of αSyn inclusions (**Fig. 1i**) Lambda scan of THK-565-stained αSyn inclusions in the M83 mouse brain indicated the peak of emission spectrum similar to the one obtained from αSyn fibrils (**Fig. 1j**). In addition, we observed that THK-565 also stained 6E10-positive amyloid-beta plaques in the arcAβ mouse brain and AT-8-positive tau inclusions in the pR5 mouse brain (**SFigs. 2a, b)**. Immunofluorescence staining of medulla oblongata brain tissue from PD patient showed detection of THK-565 in the core part of pS129-positive αSyn intraneural inclusions, Lewy bodies, and Lewy neurites (**Figs. 1k, l**).

**Fig 2.**
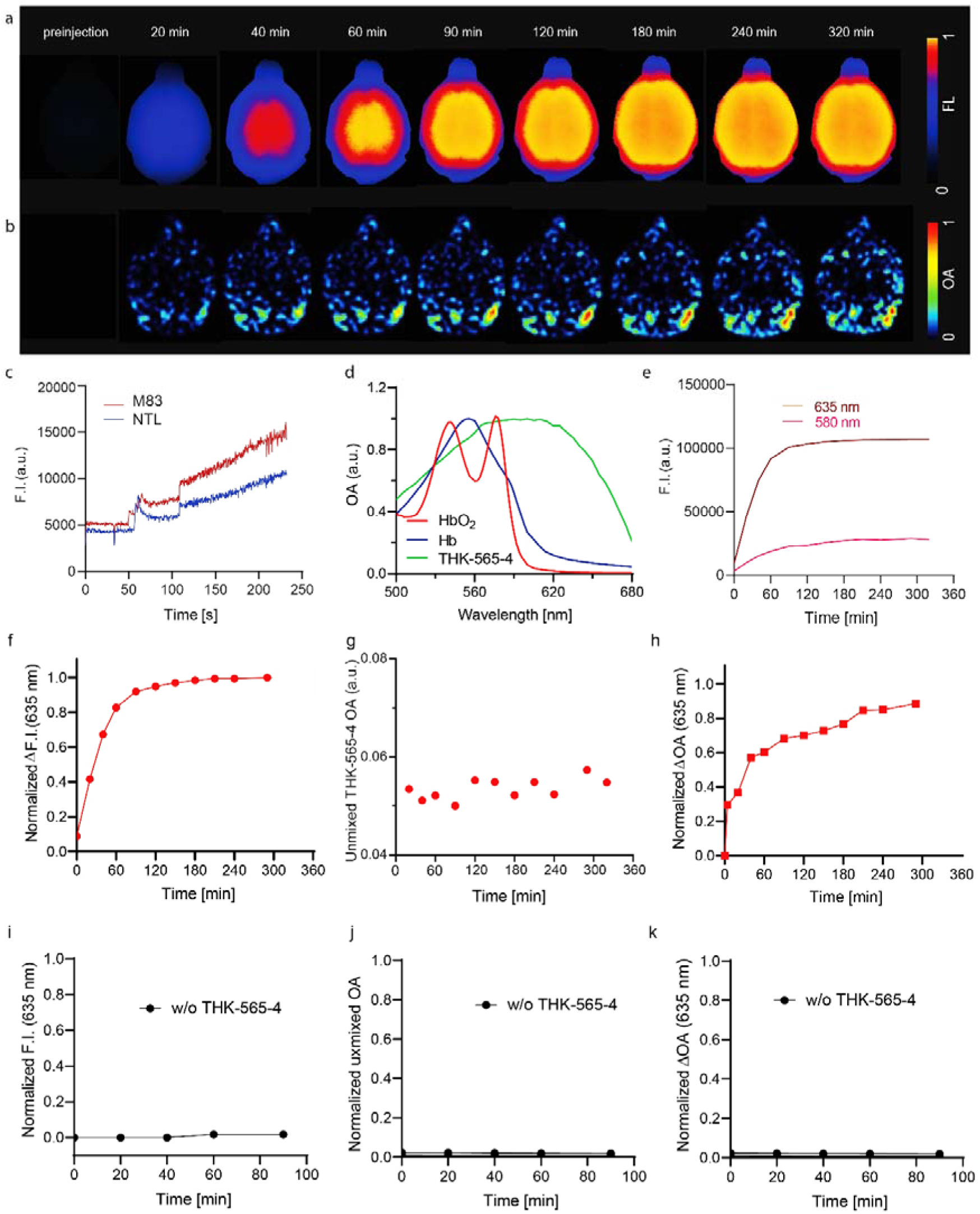
*In vivo* concurrent epifluorescence and VmsoT using THK-565. (**a**) Representative epifluorescence (FL) and vMSOT images at different time points from pre-injection of THK-565 until 320 min post-injection in the brain of one M83 mouse (horizontal view). (**b**) The time difference in the vMSOT signal during the injection of THK-565 was used to distinguish THK-565 from HbO_2_/Hb and background. (**c**) Absorbance spectrum of THK-565 (retrieved from the *in vivo* vMSOT data) and HbO_2_/Hb [41]. (**d-f**) Quantification of absolute fluorescence intensity at 580 nm and 635 nm (F. I), normalized differential fluorescence at 635 nm, normalized ΔvMSOT over the whole brain of M83 mice after THK-565 *i.v.* injection. (**g-i**) Stable normalized F.I., ΔvMSOT, and unmixed ΔvMSOT over 90 min in the brain of one M83 mouse without THK-565 injection.

### 3.2 Non-invasive *in vivo* vMSOT and epifluorescence of THK-565 uptake in the M83 mouse brain

The vMSOT imaging data analysis pipeline consisted of three steps: 3D vMSOT image reconstruction, spectral unmixing and coregistration with an MRI mouse brain atlas for VOI analysis, as described earlier [37, 45]. After *i.v.* bolus injection of THK-565, an increase in the fluorescence intensity and spectrally unmixed THK-565 signal was observed in the mouse brain parenchyma, indicating passage through the blood–brain barrier. THK-565 epifluorescence images of the brain corroborated the increase in vMOST signal associated with THK-565, albeit lacking depth information (**Figs. 2a, b**). To determine the optimal imaging time frame, we imaged one M83 mouse for 320 min post-injection of THK-565. Given that the approved *in vivo* imaging experiment was terminal, we did not keep the animal for imaging at 24 or 48 hours post-injection. The wavelengths and absorbing components were optimized so that the unmixed biodistribution of THK-565 matched the ΔvMSOT images taken at 635 nm (by subtracting a reference image obtained before injection). A clear increase/peak in the vMSOT and fluorescence signal intensity was observed upon injection, which was used for retrieving the THK-565 absorption spectrum and the unmixing analysis (**Fig. 2c**). We used five wavelengths (600, 610, 620, 630 and 635 nm) during unmixing and only HbO and THK-565 as absorbing components (**Fig. 2d**). We observed that both the fluorescence and vMSOT signals were stable at 60 min post-injection (**Figs. 2e-g**). The spectrally unmixed THK-565 appeared to have less agreement with the fluorescence dynamics compared with the differential vMSOT (ΔvMSOT) data at 635 nm (**Fig. 2h)**. A strong correlation between ΔvMSOT at 635 nm and fluorescence intensity was observed (**SFig. 3**, p < 0.0001, r = 0.9562, Pearson’s rank correlation analysis). Therefore, we used a 90 min imaging frame (20, 40, 60, 90 min) in the subsequent *in vivo* epifluorescence vMSOT experiments and relied on the ΔvMSOT signal at 635 nm for quantification [28]. To assess the stability of the fluorescence and vMSOT signal, we imaged one mouse without *i.v.* injection of contrast agent for 90 min. No alterations in the fluorescence, unmixed or differential vMSOT signal intensity over the whole brain region were observed (**Figs. 2i-k**).

**Fig. 3.**
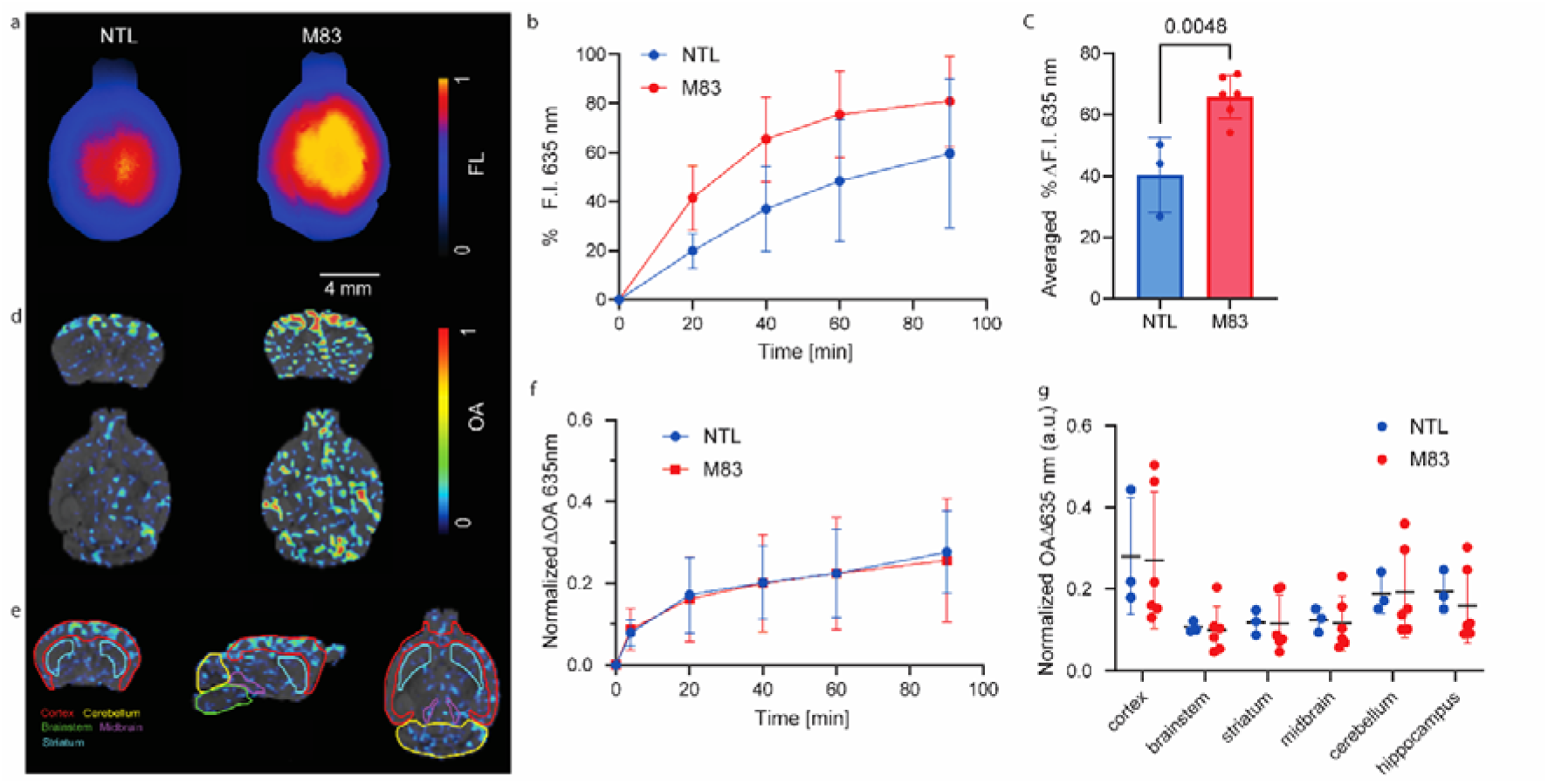
Increased THK-565 uptake in the brains of M83 mice compared to NTL mice. (**a**) Representative epifluorescence images of NTL and M83 mouse brains at 40 min post THK-565 *i.v.* injection. (**b**) % increase in fluorescence intensity over 90 min in the brains of M83 and NTL mice after THK-565 *i.v.* injection. (**c**) Higher % increase of fluorescence intensity in the brain of M83 compared to NTL mice. (**d**) Representative THK-565 signal resolved by vMSOT at 40 min post THK-565 i.v. injection (coronal and horizontal view). (**e**) Mouse brain atlas overlaid on the vMSOT images of the mouse brain (coronal, sagittal, horizontal view). (**f**) Whole brain ΔvMSOT signal intensity over 90 min of M83 and NTL mice post THK-565 *i.v.* injection. (**g**) Regional analysis of the normalized ΔvMSOT signal at 20-40 min post THK-565 *i.v.* injection.

### 3.3 THK-565 biodistribution in M83 and NTL mouse brains

M83 and NTL mice were imaged pre-, during, and post-injection of THK-565 (20 mg/kg weight, *i.v.*) using the concurrent epifluorescence-vMSOT system. Higher fluorescence intensity (635 nm) was observed in the brains of M83 mice than in those of NTL mice (**Fig. 3a**). Significantly higher percentile changes in fluorescence intensity (635 nm) were observed in the whole brain of M83 mice than in that of NTL mice (p = 0.0048, **Figs. 3b, c**). The vMSOT images acquired at 635 nm wavelength were superimposed onto the MRI atlas for VOI analysis (**Figs. 3d, e**). The differential ΔvMSOT signal corresponding to THK-565 was not significantly different in specific brain regions of M83 mice than in those of NTL mice (**Figs. 3f, g**).

### 3.4 SWI/phase and STXM imaging detect iron deposits

SWI sequences have been used to evaluate iron deposits in the brain. *Ex vivo* studies of the mouse brain using 9.4 T MRI and a cryogenic radiofrequency coil helped achieve high signal-to-noise ratios. To differentiate whether paramagnetic or diamagnetic lesions were present, a phase image was reconstructed. Iron is a ferromagnetic material and appears as black dots on both SWI and phase images. We found hypointensities in the SW images and negative phase shifts indicative of iron in the striatum of M83 mice (**Fig. 4**). Next, we validated the presence of iron by using Prussian blue staining. Blue spots in the striatum and cortex were observed in the M83 mouse brain, indicative of the presence of iron deposits (**Fig. 4**).

**Fig. 4.**
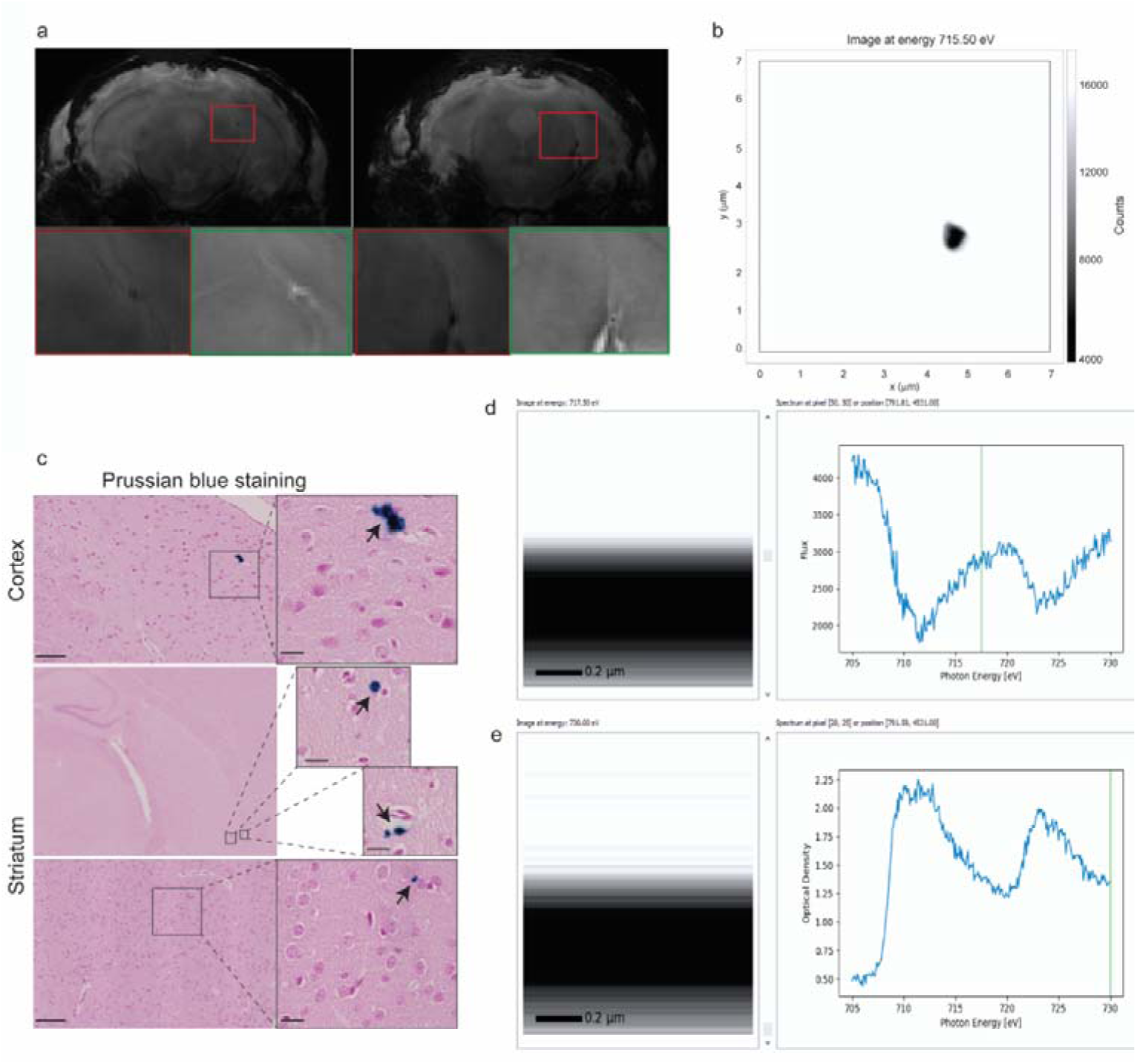
Imaging evidence of intracranial iron deposition in the M83 mouse. (**a**) Ex vivo SWI at 9.4 T and corresponding phase image showing hypointensities/negative phase shifts indicating paramagnetic iron deposition in the M83 mouse brain. (**b**) Scanning transmission X-ray microscopy showed Fe^3+^ deposits in the striatum of the adjacent brain slice of the Prussian blue-stained slice. (**c**) Prussian blue staining indicating the presence of iron deposition in the cortex and striatum of the M83 mouse brain.

An earlier study showed that ferrous and ferric iron ions show significantly different relaxation behaviours in MRI but similar susceptibility patterns [46]. Therefore, to further understand the properties of the iron deposits, we performed STXM on the adjacent brain section, in which Prussian blue staining was performed (**Fig. 4**).

### 3.5 SWI and phase imaging detect diamagnetic lesions

Calcium is a paramagnetic material, such that abnormal calcium deposits will be imaged as black dots on the SWI image (combining filtered magnitude and phase data) but as white dots on the phase image. In addition to the iron deposits, we observed hypointensities in the SW images and positive phase shifts indicative of calcification in the brains of M83 mice by *ex vivo* MRI (**Fig. 5a**). The hypointensities were more often located towards the hindbrain region below the cerebellum but also in the midbrain (**Fig. 5b**). H&E staining

**Fig. 5.**
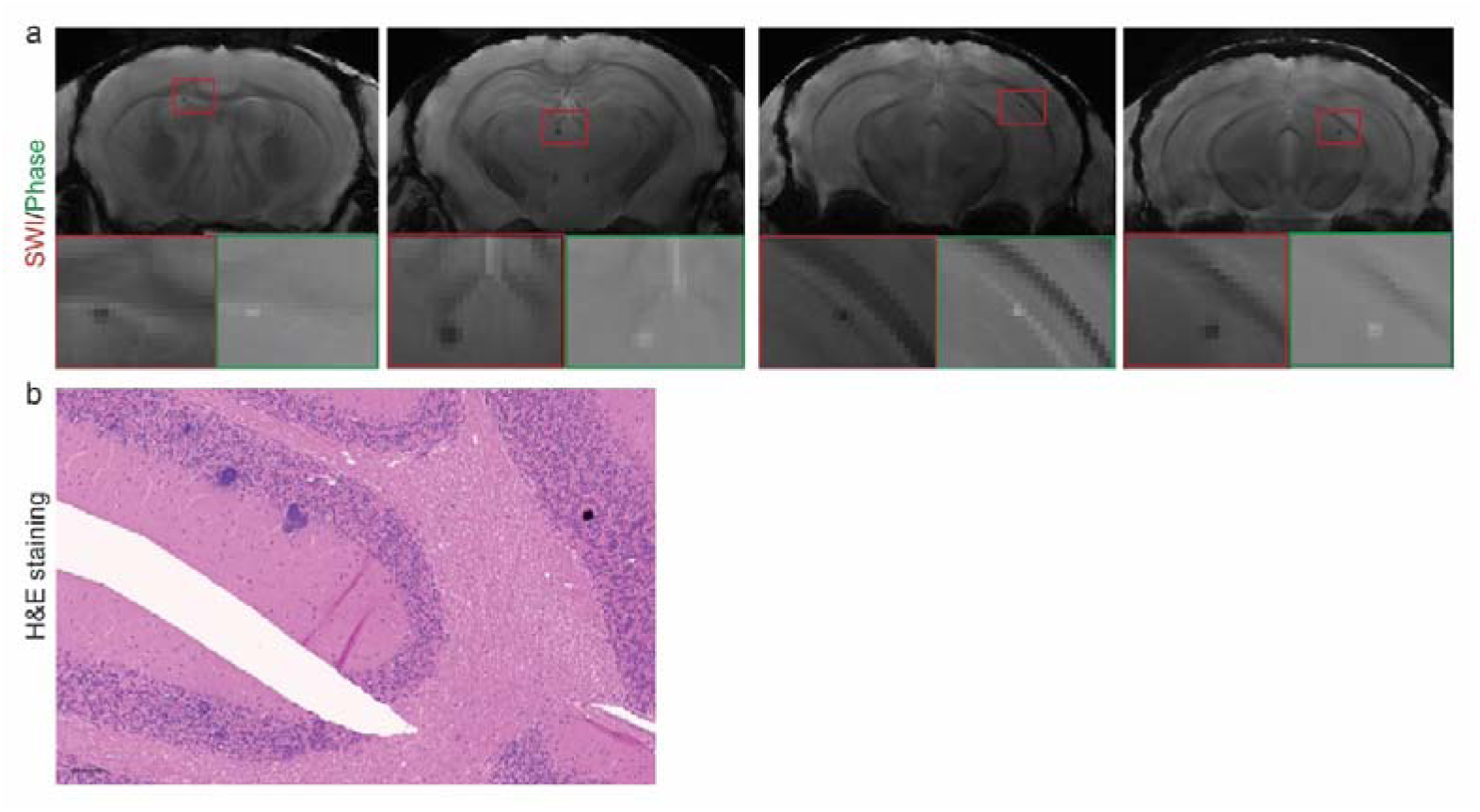
SWI and phase imaging reveal intracranial calcification in the M83 mouse. (**a**) Ex vivo SWI MRI at 9.4 T and corresponding phase image showing hypointensities/positive phase shifts indicting diamagnetic calcification in the M83 mouse brain. (**b**) H&E staining indicating the presence of calcification and enlarged perivascular space.

## 4. Discussion

New tools for non-invasive mapping of αSyn and iron deposits with high resolution facilitate understanding of disease mechanisms and the development of therapeutics [47]. Herein, we visualized the *in vivo* αSyn distribution in the brain of M83 mouse model using a hybrid epifluorescence-vMSOT system assisted by a targeted THK-565 fluorescent label. The presence of iron deposits (presumably Fe^3+^) was further demonstrated by using high-field SW MRI and STXM in the striatum and cortex of M83 mice *ex vivo*.

M83 mice show signs of motor impairment from 8 months of age. The presentation of this phenotype is associated with the formation of αS inclusion pathology throughout most of the spinal cord and brainstem, αSyn deposits start at 5 months of age and develop accumulations of αSyn in select neuronal populations, including the midbrain, cerebellum, brainstem, cortex and spinal cord [29]. The protein aggregates do not resemble Lewy bodies but are thioflavin-S-positive, indicating a fibrillar structure [29]. THK-565 facilitated sensitive detection of αSyn fibrils and inclusion slices of transgenic M83 mice and human PD brains. THK-565 was selected owing to its peak absorption at 635 nm, where light penetration is significantly enhanced with respect to shorter wavelengths. In this way, THK-565 could be more clearly distinguished from endogenous chromophores such Hb and HbO via spectral unmixing of vMSOT images acquired *in vivo*. The cortical and striatal signals detected by vMSOT *in vivo* and *ex vivo* using THK-565 are in accordance with the immunofluorescence staining results and with the reported αSyn distribution in the M83 mouse brain [29]. However, the detection sensitivity of vMSOT appeared lower than that of concurrent fluorescence recordings, which demonstrated a sufficient difference in the signal intensity between M83 and NTL mice. The lack of difference in the vMSOT signal is partly due to the large background signal fluctuations (owing to the pulsed laser instability) as well as the relatively low amount of αSyn inclusions in M83 mice at 10-11 months of age. Follow-up studies may focus on exploration of older mouse brains that were not permitted in the current animal experimentation approval.

Near-infrared fluorescence imaging detection in deep brain regions was hindered by strong absorption and scattering of the excitation light and emitted fluorescence. Submillimetre-scale intravital and multiphoton microscopies enable the visualization of αSyn deposits mainly in the cortex but are highly invasive and can only cover a very limited FOV [48–50]. Spectral unmixing can generally isolate the biodistribution of any spectrally distinctive probe from endogenous absorbers in biological tissues. However, spectral coloring effects associated with wavelength-dependent attenuation of light lead to cross-talk artifacts when considering the theoretical spectra of the absorbing substances present in the sample [51]. This is particularly important for spectral windows exhibiting sharp variations in HbO absorption, e.g., in the 605-635 nm spectral window [41].

Brain iron deposition is linked with dopamine, neuromelanin pathways, and cognitive severity in PD [52–54]. Brain iron enrichment attenuates αSyn spreading after injection of preformed fibrils [55]. Mutations in LRRK2 linked to PD sequester Rab8a to damaged lysosomes and regulate transferrin-mediated iron uptake in microglia [56]. A few MRI studies have been performed in animal models of PD, such as by using T_2_* [3, 4, 57], as well as in patients with PD using novel MRI contrasts, SWI, quantitative susceptibility mapping, and R_2_ R_2_* [5, 6, 8, 58–63]. A previous study using 3D elemental bioimaging showed Fe, Zn, Cu, Mn and P in a 6-hydroxydopamine-lesioned mouse brain [64, 65]. Here, we demonstrated regional hypointensities in the hippocampus, cortex, striatum, midbrain and thalamus by using SW images of M83 mice, which in corresponding phase images indicated paramagnetic (iron) lesions in the brain of M83 mice. Moreover, the STXM detection of Fe^3+^ deposits in the adjacent slice of Prussian blue-stained iron deposits in the striatum further validated this finding. The striatum is a vulnerable brain region that loses its dopaminergic innervation and is affected early in PD patients and transgenic animal models of PD [66]. The striatum and cortex of the M83 mouse thus bear both αSyn inclusions and iron deposits. In addition, we also observed the presence of calcification in the M83 mouse brain by SW/phase MR. Basal ganglia calcifications [67] have been reported in patients with PD [68].

There are several limitations in the current study. First, vMSOT imaging data are subject to spectral coloring due to wavelength-dependent light absorption in biological tissue. However, reliable fluence correction is still an unsolved problem; thus, we opt not to apply complex processing [69]. Second, longitudinal studies are required to determine the sensitivity and specificity of the proposed methodology, at which age of M83 mice can THK-565-positive αSyn inclusions be detected by vMSOT, and whether it can follow the spreading of αSyn in the brain. Given the relatively low load of αSyn inclusions in the transgenic animal model at this age, αSyn preformed fibril-injected animals or AAVαSyn models might have an advantage in the assessment of αSyn imaging tracers with the possibility of a high load of αSyn inclusion in the brain. On the other hand, later disease stages at 18-24 months of age in M83 mice also shed light on the ability of these methodologies to monitor disease progression. Such studies in aged M83 mice were limited by a lack of local veterinary approval due to phenotypic constraints.

In conclusion, we demonstrated successful non-invasive imaging of αSyn in M83 mice with a concurrent epi-fluorescence-vMSOT system at ∼110 mm spatial resolution. This *in vivo* imaging platform provides a new tool to map αSyn distribution in αSyn mouse models, which may facilitate the monitoring of αSyn-targeting therapeutics.

## Acknowledgement

The authors acknowledge Dr. Saroj Kumar Rout at ETH Zurich, Mr. Michael Reiss, Prof. Jan Klohs, Ms. Agathe Tournant at the Institute for Biomedical Engineering, ETH Zurich/University of Zurich, Ms Man Hoi Law at Imperial Colleague London; Mr Vasil Kecheliev, Mr Daniel Schuppli at the Institute for Regenerative Medicine, University of Zurich; and ZMB for technical assistance.

## Author Contributions

The study was designed by RN. NO provided THK-565. JG performed recombinant fibril production and binding measurements. ZC and DR designed and built the hybrid fluorescence and vMSOT system. XLDB, NS, and RN performed *in vivo* imaging. ID and DN performed animal breeding and genotyping, AT, ML, and RN performed *ex vivo* MRI. RS, LJ, and IR performed thin sample preparation and scanning X-ray microscopy. BC and DN performed histology and microscopy. NS, BC, XLDB, and RN performed the data analysis. NS, BC, DN, JG, XLDB, NO, DR, RN interpreted the data. RN wrote the first draft. All authors contributed to the revision of the manuscript. All authors read and approved the final manuscript.

## Disclosures

The authors declare no conflicts of interest.

### Consent to participate

Not applicable.

### Consent for publication

Not applicable.

## Supplementary files

**Supplementary Fig 1.**
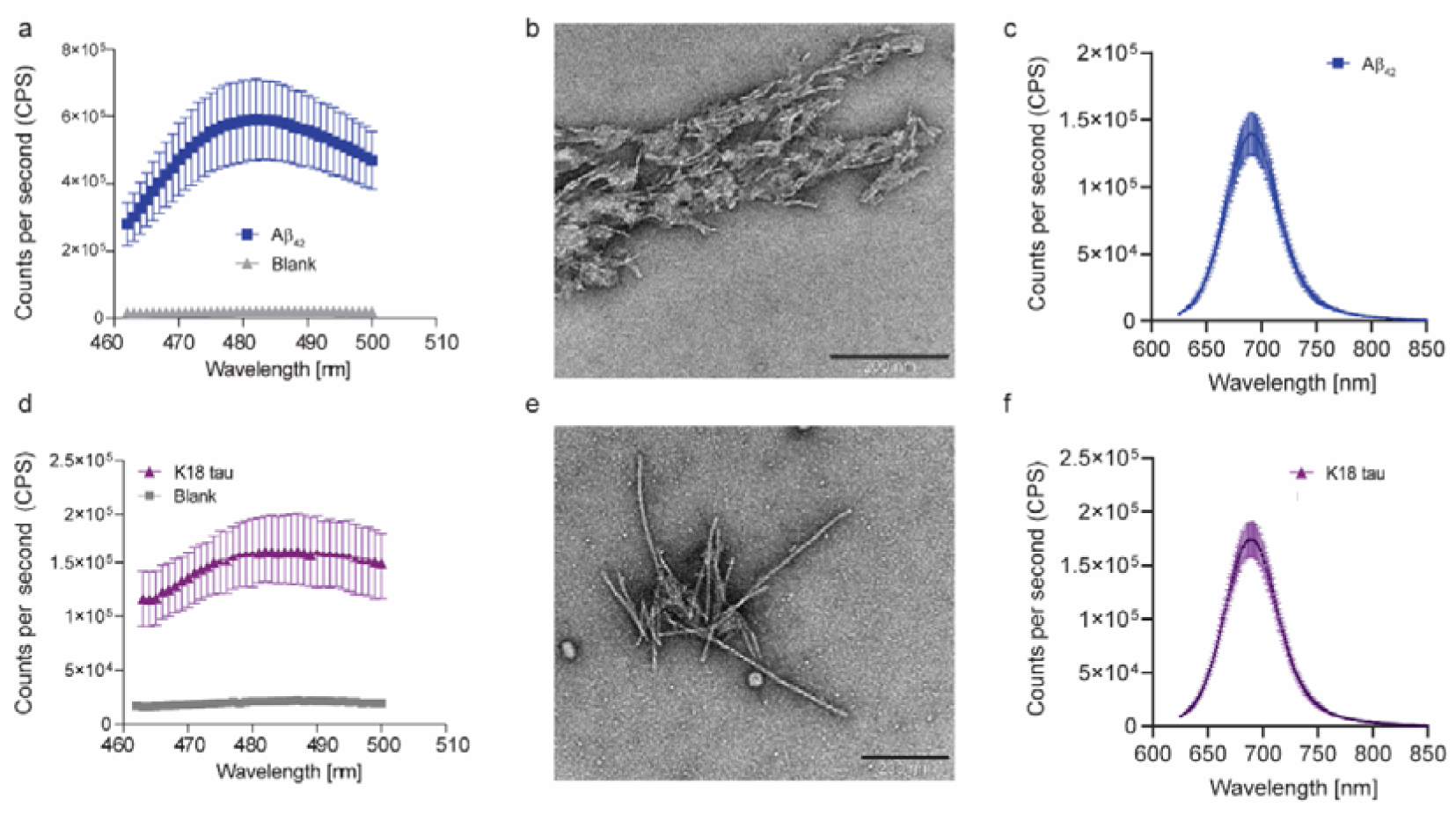
Characterization of THK-565 on recombinant Aβ_42_ and K18 tau fibrils. (**a, d**) Thioflavin T assay of recombinant Aβ_42_ (blue) and K18 tau (purple) fibrils and blank (gray); (**b, e**) Transmission electron microscopy characterization of recombinant Aβ_42_ and K18 tau fibrils. Scale bar = 200 nm; (**c, f**) Spectrofluorometric measurements of the binding of THK-565 to recombinant Aβ_42_ and K18 tau fibrils.

**Supplementary Fig 2.**
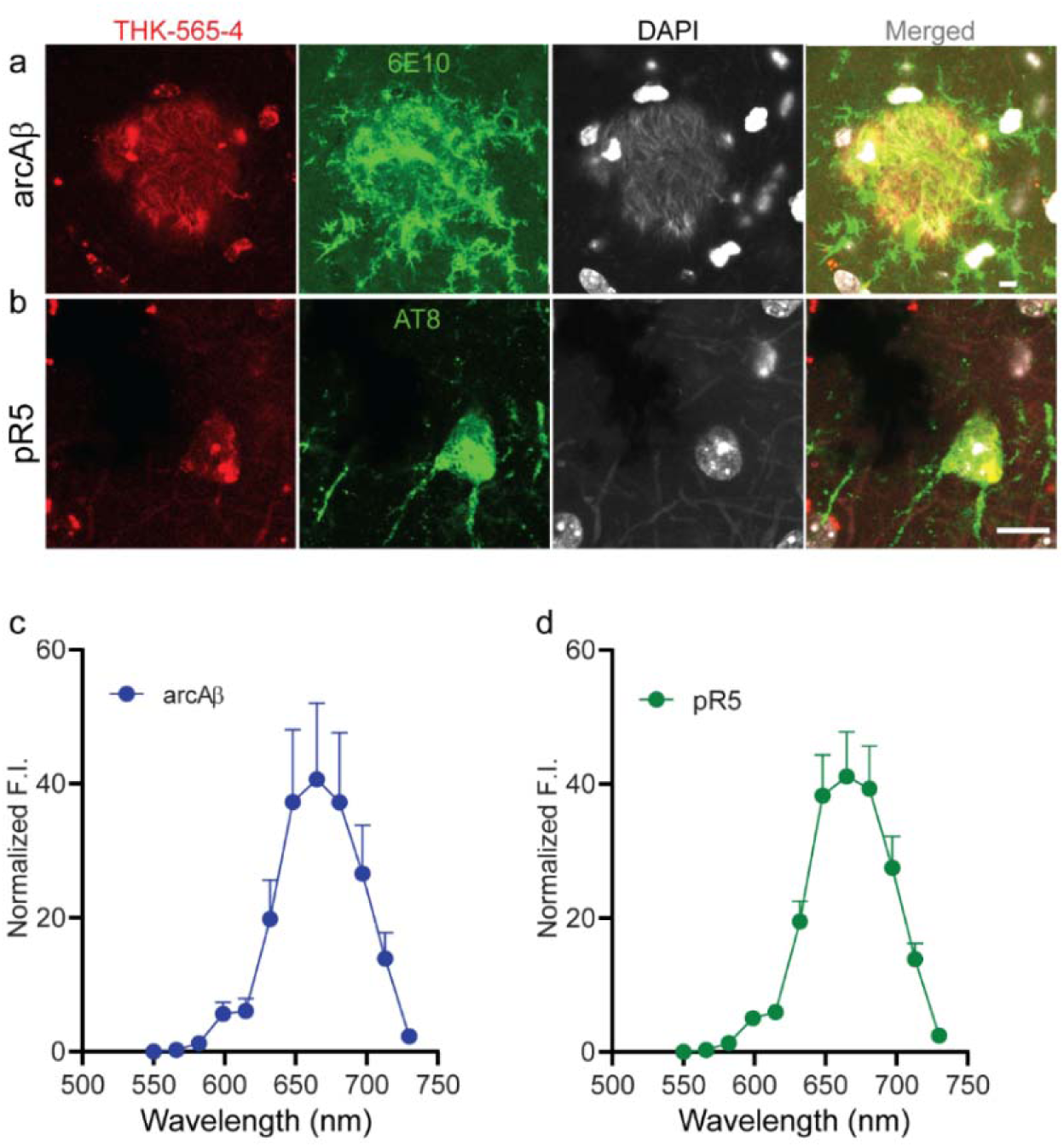
Staining of THK-565 in amyloid-beta plaques and tau inclusions in arcAβ and pR5 mouse brains. (**a, b**) Immunofluorescence staining using THK-565 (red), with anti-Aβ antibody 6E10 (green) on arcAβ mouse brain and anti-phospho-tau antibody AT-8 (green) on pR5 mouse brain; nuclei were counterstained using DAPI (gray). Scale bar = 10 μm; (**c, d**) Lambda scan of THK-565-stained Aβ plaques and tau inclusions on arcAβ and pR5 mouse brains using confocal microscopy.

**Supplementary Fig 3.**
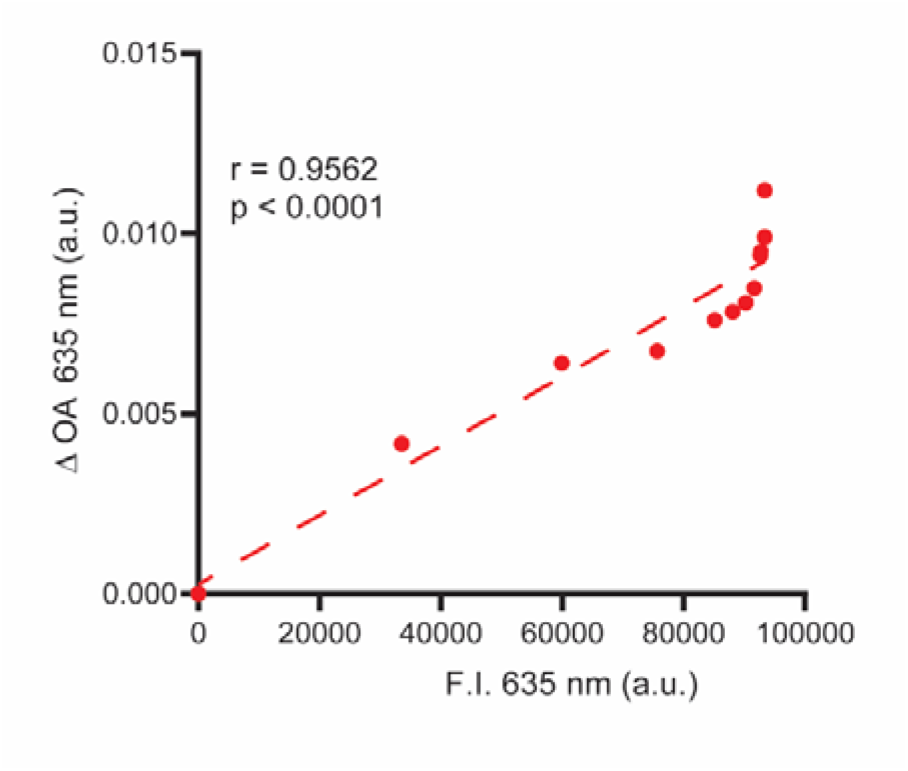
There was a strong correlation between the ΔvMSOT signal acquired at 635 nm and fluorescence intensity. Imaging in a representative M83 mouse after *i.v.* injection of THK-565; Pearson’s correlation.

**Supplementary Table 1.** List of primary antibodies used for western blotting and immunofluorescence staining.

| Antibody/Compounds | Company | Cat. No. | Dilution |
| --- | --- | --- | --- |
| DAPI | Sigma□Aldrich | D9542-10MG | 1:1000 |
| Goat-anti-Rabbit Alexa488 | Invitrogen | A11034 | 1:200 |
| 6E10, Anti-β-amyloid, 1-16 antibody | Biolegend | 803001 | 1:1000 |
| AT-8, anti-phospho-tau Ser202, Thr205 | Invitrogen | MN1020 | 1:1000 |
| Syn303, anti-α-Synuclein (aa 1–5) | Biolegend | MMS-5085 | 1:1000 |
| LB509, anti-α-Synuclein (aa 115-122) | EMD Millipore | MABN824 | 1:1000 |
| pS129 (phospho-Ser129) | Abcam | ab51253 | 1:1000 |
| VECTASHIELD® Antifade | Vector Laboratories | H-1000-10 |  |
| Normal goat Serum (NGS) | Chemie Brunschwig | JAC005-000-121 |  |
| Normal Donkey Serum (NDS) | Southern Biotech |  |  |
| Triton X-100 | Sigma□Aldrich |  |  |
| Anti-Tau (RD4) Antibody, clone 1E1/A6 | Sigma□Aldrich | 05-804 | 1:1000 |
| Anti-α-Synuclein mouse monoclonal antibody, clone Syn211 | Sigma□Aldrich | S5566 | 1:2000 |
| Monoclonal Anti-β-amyloid antibody, clone BAM-10 | Sigma□Aldrich | A5213 | 1:1000 |
| Anti-mouse IgG antibody conjugated to horseradish peroxidase (HRP) | Jackson ImmunoResearch, | 115-035-166 | 1:5000 |
| THK-565 | NO |  | 16.3 |
| ECL Prime Western Blotting Detection Reagents | Cytiva | RPN2232 |  |
| Fomblin Y, LVAC 16/6, average molecular weight 2700 | Merck | 317942 |  |

